# Tuning in on auditory details is difficult: Individuals with aphasia show impaired acoustic and phonemic processing

**DOI:** 10.1101/2022.12.14.520503

**Authors:** Jill Kries, Pieter De Clercq, Robin Lemmens, Tom Francart, Maaike Vandermosten

**Affiliations:** Experimental Oto-Rhino-Laryngology, Department of Neurosciences, Leuven Brain Institute, KU Leuven, Leuven, Belgium; Experimental Neurology, Department of Neurosciences, KU Leuven, Leuven, Belgium; Laboratory of Neurobiology, VIB-KU Leuven Center for Brain & Disease Research, Leuven, Belgium; University Hospitals Leuven, Department of Neurology, Leuven, Belgium

## Abstract

Acoustic and phonemic processing are understudied in aphasia, a language disorder that can affect different levels and modalities of language processing. For successful speech comprehension, processing of the speech envelope is necessary, which relates to amplitude changes over time (e.g., the rise times). Moreover, to identify speech sounds (i.e., phonemes), efficient processing of spectro-temporal changes as reflected in formant transitions is essential. Given the lack of aphasia studies on these aspects, we tested rise time processing and phoneme identification in 29 individuals with post-stroke aphasia and 23 healthy age-matched controls. We found significantly lower performance in the aphasia group than in the control group on both tasks, even when controlling for individual differences in hearing levels and cognitive functioning. Further, by conducting an individual deviance analysis, we found a low-level acoustic or phonemic processing impairment in 76% of individuals with aphasia. Additionally, we investigated whether this impairment would propagate to higher-level language processing and found that rise time processing predicts phonological processing performance in individuals with aphasia. These findings show that it is important to develop diagnostic and treatment tools that target low-level language processing mechanisms.

## 1 Introduction

Aphasia is an acquired language disorder that frequently occurs after a cerebrovascular accident (CVA), or stroke. Given that a stroke can impact diverse brain areas to a varying amount, the symptoms and severity of aphasia are heterogeneous, encompassing difficulties across all speech processing levels, e.g., acoustic, phonological, semantic or syntactic, and language modalities, e.g., speech comprehension, production, reading or writing [1, 2]. Aphasia research mostly covers the assessment of higher-level language functions such as phonology, semantics or syntax [2, 3]. Behavioral tests of lower-level comprehension functions, such as auditory spectro-temporal processing, are not part of the clinical test protocol [4, 5, 6] and these aspects have been researched rather sparsely [7, 8]. In particular, spectro-temporal processing of acoustic (e.g., dynamic amplitude changes, such as rise times) and phonemic (e.g., dynamic frequency changes that help identifying phonemes) aspects have not yet been assessed in individuals with aphasia (IWA), even though they are crucial for speech understanding [9, 10, 11, 12]. Only one study explored spectro-temporally modulated sounds in individuals with severe Wernicke’s aphasia [8]. Interestingly, they found impaired dynamic auditory processing in their sample, which correlated with comprehension and phonological tests [8]. Similarly, research in the field of developmental dyslexia demonstrated a link between tasks measuring dynamic acoustic and phonemic processing and higher-level phonological processing [13, 14, 15, 16, 17, 18]. Therefore, we investigated (1) whether a significant difference can be found between IWA and healthy controls based on rise time processing and phoneme identification tasks and (2) whether these tasks can predict phonological processing performance in IWA.

### 1.1 Rise time processing

Auditory processing of dynamic changes in the amplitude of speech is crucial for segmenting speech into meaningful sublexical and lexical units and thus, for speech comprehension [11, 12, 19, 20]. In fact, one of the most important auditory cues for speech comprehension is the envelope, i.e., amplitude changes over time [9, 12]. Shannon et al. [9] found that listeners can understand speech based on the temporal envelope alone. Further, Oganian and Chang [12] investigated what landmark of the envelope would be encoded strongest in the superior temporal gyrus and demonstrated that it is the rate of amplitude change at acoustic onsets of the envelope, rather than the absolute amplitude. Processing of the rate of positive change (rise) in amplitude, here referred to as rise time, can be measured via a rise time discrimination (RTD) task. Efficient processing of rise time is important for identifying the onset of a phoneme or a syllable and hence, aids speech segmentation. Furthermore, the rise time gives information about the syllable stress, since stressed syllables have steeper rise times than unstressed ones [12, 19, 20]. Hence, rise time processing is crucial for comprehension.

Biedermann et al. [21] observed that patients with a lesion in the right or left auditory cortex (due to middle cerebral artery damage), as well as patients with a lesion in the subcortical auditory structures, showed auditory processing impairments in signal discrimination tasks in noise targeting frequency, amplitude and duration aspects. The middle cerebral artery supplies the insular and auditory cortex (among other areas) with oxygenated blood [22]. An estimated 70% of stroke-induced aphasia cases result from a stroke that involves the middle cerebral artery to some extent [23, 24], hence an auditory processing impairment in a considerable number of IWA can be expected. Despite this fact and the importance of efficient dynamic acoustic processing of the amplitude for speech comprehension, the aphasia literature has mostly neglected studying this feature of speech processing. However, some studies in IWA have focused on other, non-dynamic acoustic cues, such as gap or stimulus duration [7, 25, 26, 27]. These studies found impaired processing of gap and stimulus duration changes in IWA compared to healthy controls in a variety of tasks, i.e., stimulus (tone or vowel) duration judgement tasks [25, 26], temporal order detection tasks based on gap duration changes [7] and gap detection tasks [26, 27].

Assessment of RTD performance in children and adults with developmental dyslexia has shown that they performed significantly lower than healthy controls [28, 29]. This task can thus be seen as a behavioral marker of dynamic acoustic processing deficits that seem to underlie phonological problems in individuals with reading difficulties (i.e., developmental dyslexia). In the present study, we investigate whether RTD performance can also present a behavioral marker of dynamic acoustic processing impairments in individuals with post-stroke aphasia.

### 1.2 Phoneme identification

Although efficient rise time processing supports identification of speech sounds (i.e., phonemes), efficient processing of dynamic changes in frequency is also required. In order to identify phonemes regardless of speaker-related acoustic variability, e.g., variation in accent, speed or syllable stress, certain acoustic cues need to be inhibited to identify the correct phoneme category [30]. Phoneme identification is often overlooked in aphasia assessments. Especially spectro-temporal aspects of phoneme identification, such as processing of formant transitions (i.e., spectro-temporal changes in the speech signal defined by vocal tract movements), have not yet been investigated in IWA. Phoneme identification can be assessed by presenting ambiguous speech sounds between two similar phonemes, e.g., /bA/ and /dA/ [31]. This task offers a measure of how consistently speech sounds with some acoustic variation are classified into the same category and thus, how clearly defined the borders of the phonemic representations are [30, 31].

Functional magnetic resonance imaging (fMRI) research has shown that phoneme identification is localized in the left superior temporal gyrus and sulcus [32, 33, 34]. In IWA after stroke, it is not uncommon that the lesion coincides with this area supplied mostly by the middle cerebral artery [21, 23]. Accordingly, Robson et al. [8] found impaired processing of sound stimuli with sinusoidal spectro-temporal modulations in individuals with severe Wernicke’s aphasia, who at group-level showed the largest lesion overlap in the temporo-parietal junction and temporal lobe. The type of spectro-temporal modulations used for their stimuli [8] are however different from formant transitions, necessary for phoneme identification. Another study by Fink et al. [7] reported lower performance on a phoneme discrimination task at word-level in IWA compared to a control group. However, the phoneme contrast used in the study (/d/ versus /t) by Fink et al. [7] purely relied on temporal differences, i.e. differences in voice onset time, and did not assess processing of dynamic spectro-temporal cues (formant transitions), which are important for phoneme identification. Given the involvement of this mechanism for speech comprehension, information about the consistency of identifying speech sounds into the same phonemic category may be important for diagnosis and therapy of speech processing problems in aphasia.

### 1.3 Association between acoustic-phonemic processing and higher-level speech processing

Assessing acoustic and phonemic processing to inform therapy plans would be particularly helpful if a relation with higher-level speech processing performance would be present in aphasia. Neural models show that different speech processing mechanisms overlap in time and that the lower-level auditory analysis interacts bidirectionally with top-down contextual information to aid quick access to semantic representations [35, 36]. Hence, the interaction between these systems might be adversely affected if one of the systems is defective. To explore the link between lower- and higher-level mechanisms, it has been investigated whether an impairment at lower-level speech processing would affect higher-level speech processing. In the field of developmental dyslexia, evidence suggests that impairments in auditory processing (measured via psychoacoustic tasks) propagate onto higher-level mechanisms, such as phonological processing [37]. Specifically, Boets et al. [18] found a link between categorical perception of phonemes and phonological awareness in kindergarteners at familial risk for dyslexia. Furthermore, rise time processing predicted phonological processing performance and literacy measures in individuals with dyslexia [13, 14, 15, 16, 17].

In IWA, auditory discrimination of stationary stimuli has been linked to phonological processing performance [7, 8, 26]. Furthermore, Robson et al. [8] also found a link between dynamic auditory processing and comprehension and phonological tests in IWA. To date, it is however not yet clear whether rise time processing and phoneme identification are also associated with phonological processing in IWA. If so, then targeting these acoustic and phonemic cues in interventions for aphasia could potentially result in a cascading effect on higher-level linguistic processing aspects.

### 1.4 Current study

Our primary aim was to explore acoustic and phonemic processing in IWA and a control group by administering the RTD task and the phoneme identification task. The RTD task is a measure of (non-linguistic) auditory processing targeting dynamic changes in amplitude. The phoneme identification task indicates how well the phoneme boundaries are defined, based on subtle spectro-temporal changes in syllable-level stimuli. Both tasks are presented auditorily and thus require functional peripheral hearing, which is often impaired in older subjects, such as the ones included in our study. We therefore statistically controlled for the influence of hearing levels. Furthermore, the two tasks also involve cognitive processes, such as attention and executive functioning. IWA frequently show concomitant cognitive impairments and thus, we also statistically controlled for the influence of cognitive functioning. We hypothesized IWA to show lower performance at group-level than the control group on the RTD and the phoneme identification task. To explore how many IWA would deviate from the control group, hence display impaired performance on the acoustic and phonemic tasks, we implemented an individual deviance analysis [38]. The second aim was to investigate whether the performance at the acoustic and phonemic tasks would predict performance at higher-level phonological tasks within the aphasia group. We expected scores on the acoustic-phonemic tasks to predict performance on the phonological processing tests.

## 2 Methods

We compared acoustic and phonemic processing in IWA and age-matched healthy controls via two pyschoacoustic tasks, i.e., the RTD task and the phoneme identification task. To explore whether acoustic and phonemic processing skills in IWA are associated with higher-level phonological impairments, we also administered two language tests that measure phonological processing. Furthermore, a validated picture-naming task and a general aphasia test were administered to characterize the aphasia sample.

### 2.1 Participants

We tested 29 IWA in the chronic phase (≥ 6 months) after stroke (time since stroke in months: mean=38.8, standard deviation (SD)=70.7, median=18, min=6, max=368) and 23 healthy age-matched controls. All participants were Dutch native speakers from Flanders, Belgium. IWA were recruited in two ways. Between October 2018 and March 2021 (with a COVID-19-related break between March and June 2020), patients at the stroke unit of the university hospital (UZ Leuven) were systematically screened for language deficits on a daily basis using the Language Screening Test (LAST) [39]. For this, they had to give informed consent according to the declaration of Helsinki. The study received ethical approval by the medical ethical committee of KU Leuven and UZ Leuven. Patients with a stroke that scored equal to or below the cut-off score were contacted earliest 6 months after the stroke to participate in the study. They also had to meet further inclusion criteria before they were contacted, i.e., having no formal diagnosis of a psychiatric or neurodegenerative disorder, and having a left-hemispheric or bilateral lesion. The second recruitment strategy for IWA encompassed contacting independent speech-language pathologists (SLP) and rehabilitation centra in Flanders to advertise the study via flyers and posters (see fig. S.1 for a flowchart of the recruitment strategies). Healthy age-matched controls (n=23) were recruited via flyers positioned in recreational community centers for elderly. The accepted age for participation of healthy controls was gradually adapted based on the mean age and SD of IWA included in the study.

The resulting aphasia sample from the two recruitment strategies was checked for language impairments using two standardized diagnostic aphasia tests, i.e., the ScreeLing [40] and the Dutch naming test (Nederlandse Benoem Test (NBT)) [41]. The ScreeLing test does not include a picture-naming task, therefore we added the NBT to characterize aphasia. The NBT is a validated picture-naming test with 92 items and a maximum score of 276 points (cut-off threshold: 255) [41]. The ScreeLing is a validated test for diagnosis and therapy follow-up that consists of three subtests, i.e., phonology, semantics and syntax, each containing four tasks with 24 items [40]. Here, we administered the ScreeLing on a tablet using the Gorilla Experiment Builder (http://www.gorilla.sc) [42]. To check language impairments among IWA, we used the total score of the ScreeLing (maximum score: 72, cut-off threshold: 68). The test scores of IWA can be found in table 1 and they are visualized in supplementary figure S.2. We included individuals that scored either (1) below the cut-off threshold on at least one of these two tests at the moment of data collection (n=20) (table 1), or (2) had a documented language impairment in the acute phase (8 of the 9 remaining individuals scored below cut-off threshold in the acute phase on the ScreeLing (n=5), the Comprehensive Aphasia Test-NL (CAT-NL) (n=2) or the Aachen Aphasia Test (n=1) and one IWA, who was referred to the study via flyer, provided medical proof of a diagnosis of severe motor aphasia in the acute phase). Also note that 8 out of the 9 IWA, who did not score below the cut-off thresholds on neither the ScreeLing nor the NBT at the time of data collection, were still following speech-language therapy at the time of data collection (table 1). All participants (or their legal guardian) gave informed consent prior to participation and this part of the study also got ethical approval by the the medical ethical committee of KU Leuven and UZ Leuven.

**Table 1:**
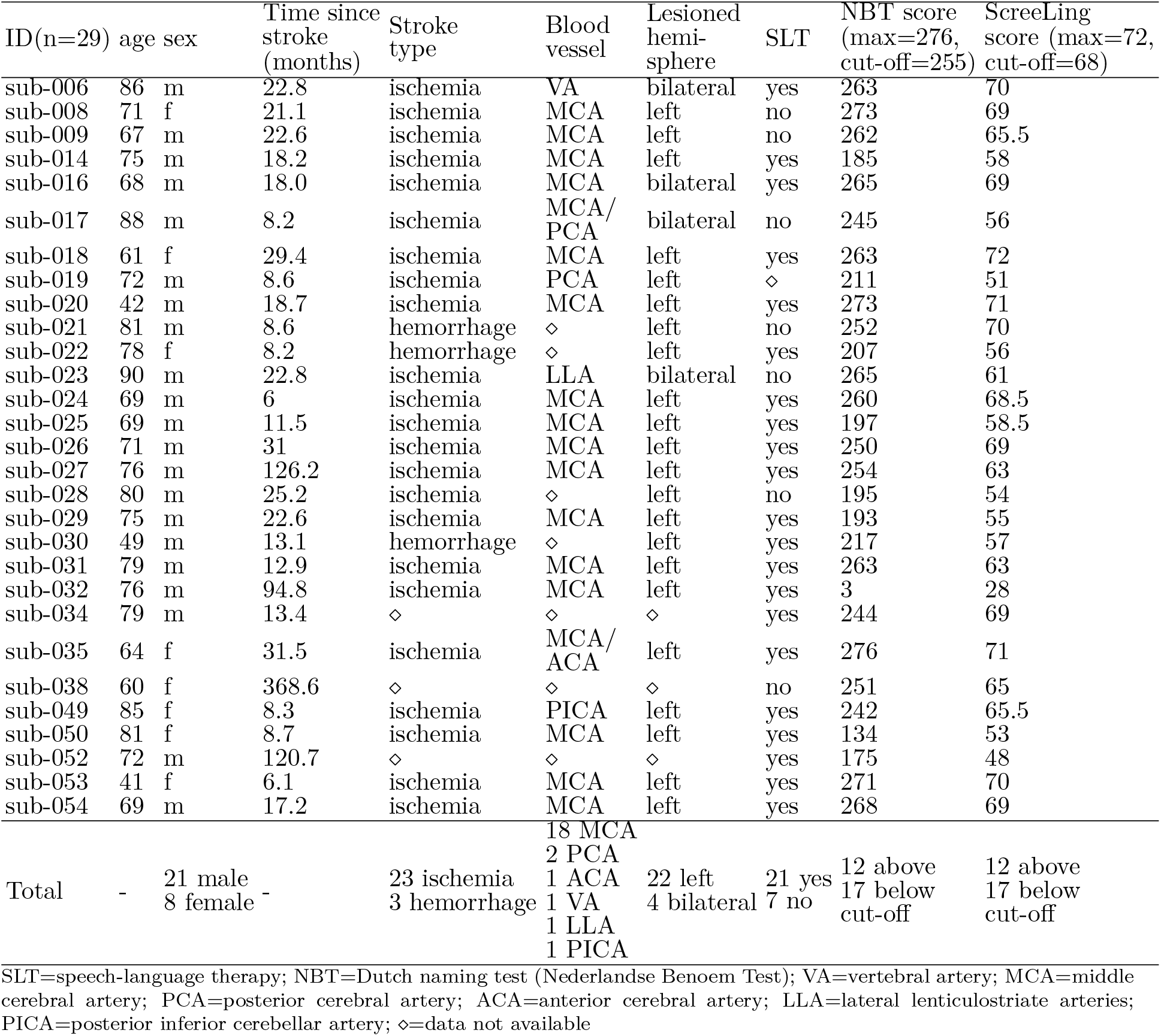
Demographics and lesion information of the aphasia group.

IWA were on average 71.52 years old (SD: 12.15) and controls were 71.52 years old (SD: 7.15). No age difference was found between groups (W= 365.5, p= 0.56) (supplementary fig. S.2). The sex ratio was not significantly different between both groups (*χ*^2^= 3.4e-31, df= 1, p= 1; IWA: 27.6% female, 72.4% male; controls: 30.4% female, 69.6% male). The level of education did also not differ between groups (*χ*^2^= 5.101, df= 4, p= 0.277; supplementary table S.1). The groups significantly differed on the NBT (W= 72, p= <0.001) and on the ScreeLing (W = 121, p= <0.001), as we expected given the inclusion criteria (supplementary fig. S.2). These variables and more demographic information (time since stroke, speech-language therapy) as well as lesion information (stroke type, blood vessel blocked or ruptured, lesioned hemisphere) about the aphasia sample can be found in table 1. Note that out of the IWA of whom we had access to lesion information, 81.8% had a lesion in the middle cerebral artery.

### 2.2 Behavioral measures used for statistical analyses

#### 2.2.1 Rise time discrimination task

The RTD task measures how well participants discriminate the rate of change in amplitude at the onset of a sound. Precisely, the task was presented as a three-alternative forced choice task, where the deviant stimulus had to be discriminated from two identical reference stimuli (fig. 1A and B). The stimuli were created in MATLAB [43] using one-octave noise bands centered at 1 kHz [29]. The software APEX was used to present the task [44]. Stimuli were calibrated and presented in the left ear at 70 dB SPL. The reference stimulus had a rise time of 15 milliseconds (ms). The deviant stimuli were computed to have rise times that decreased logarithmically in 50 steps from 699 ms to 16 ms. The duration of each stimulus was 800 ms. The number of trials differed between participants, as the task followed a one-up/two-down adaptive staircase procedure. This means that after two correct responses in a row, the difference in rise time between stimuli became smaller, thus more difficult, during the next trial. After one erroneous response, the difference in rise time between stimuli became larger, thus easier to discriminate. This way, a threshold corresponding to 70.7% correct was targeted [45]. The task ended once 8 reversals (i.e., changes in direction) were reached. In case no reversals were present, the task ended after a maximum of 87 trials. The individual performance trajectories as well as the group average and standard error (SE) of the trajectories are visualized in figure 2A. The rise times of the deviant stimulus of the last 4 reversals were averaged to determine the final threshold. This threshold was used for statistical analyses.

**Figure 1:**
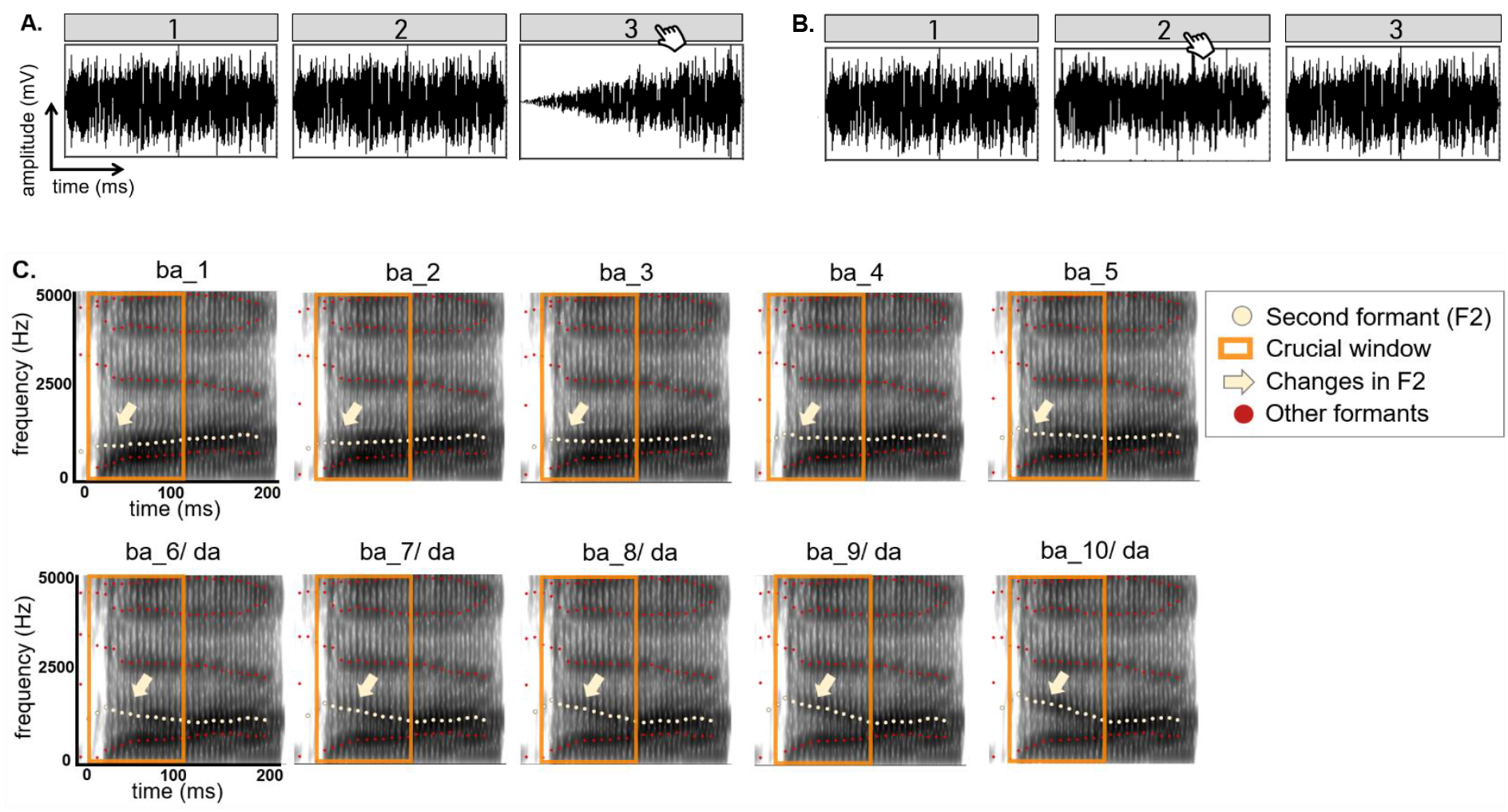
Stimuli used for the auditory-phonemic tasks. **A.** Visualization of the RTD task with an example of the largest rise time contrast between the reference stimuli (15 ms rise time) and the deviant stimulus (699 ms rise time). The deviant stimulus position was randomized across trials. **B.** Visualization of the RTD task with an example of the smallest rise time contrast between the reference stimuli (15 ms rise time) and the deviant stimulus (28 ms rise time) identified by the highest performing participant in this study. In A. and B. the deviant stimulus is indicated by the hand cursor. **C.** Visualization of the 10 stimulus steps used for the categorical perception task. The yellow dots depict F2, the yellow arrow points to the change in the slope within the first 100 ms of each stimulus (orange square). After 100 ms, the stimuli remain unchanged. The other formants, depicted by the red dots, also remain unaltered across stimulus steps.

**Figure 2:**
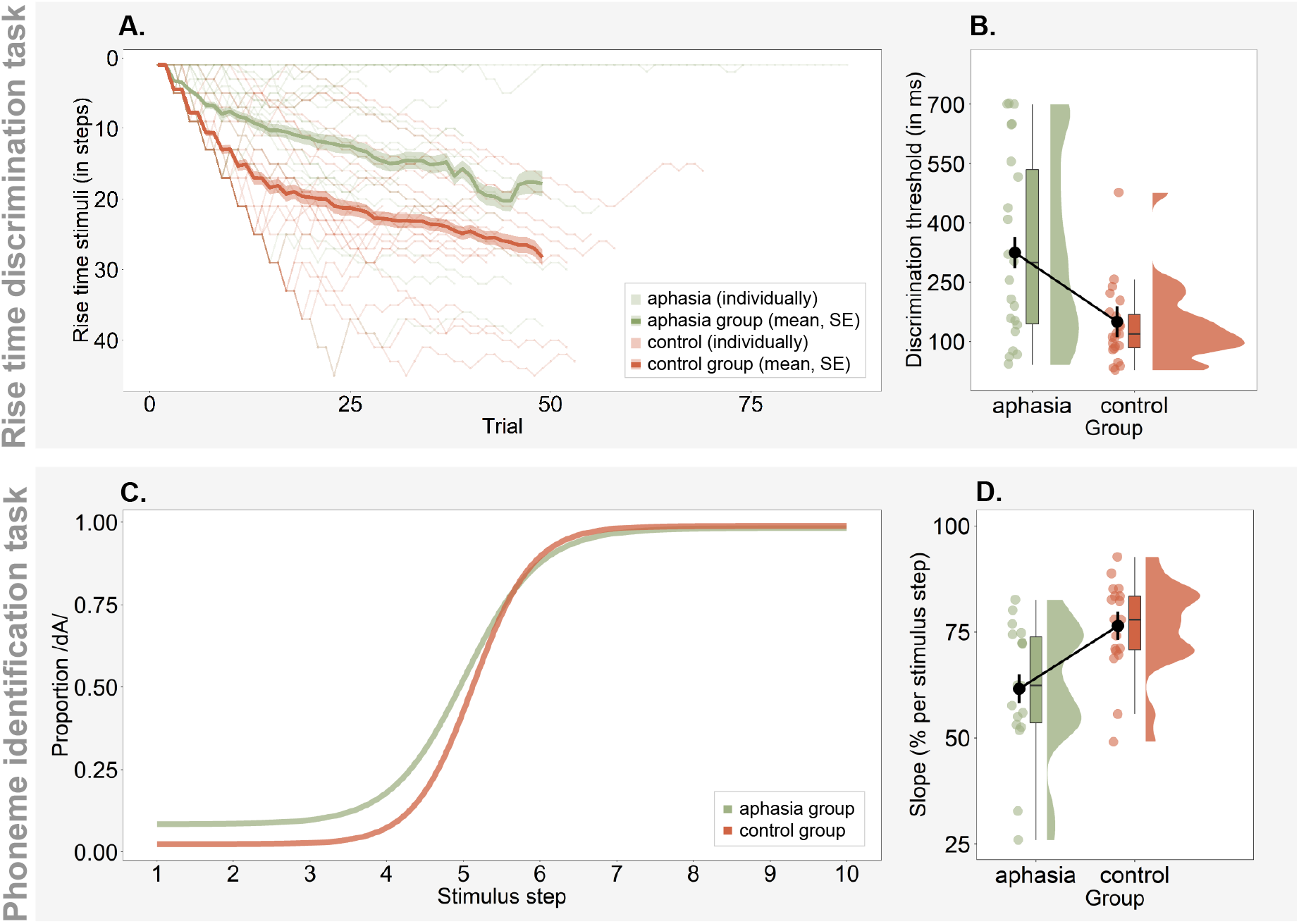
IWA show decreased performance at two auditory-phonemic processing tasks. **A.** The RTD thresholds were determined via a one-up/two-down adaptive staircase procedure. This figure shows the individual performance trajectories as well as the group average and standard error across trials for visualization purpose. The deviant stimulus rise times of the last 4 reversals were averaged to determine the final threshold, which was used for statistical analysis. **B. IWA** at group level processed the rate of change in amplitude less precisely (i.e., larger differences in rise time are needed to discriminate stimuli) than healthy, age-matched controls, even when controlling for hearing and cognition. **C.** Visualization of the average of subject-specific psychometric functions by group, representing performance on the phoneme identification task. The steeper the slope, the more consistently a phoneme is identified, hence suggesting better phoneme representation. **D. IWA** showed significantly lower slopes than age-matched controls at group level on the phoneme identification task, even when controlling for hearing and cognition.

To make sure that the task was well understood by all participants, they performed between 4 and 8 practice trials before starting the task. In the aphasia group, 23/29 IWA completed RTD task, while 6 IWA experienced the task as too difficult after the initial trials. All 23 healthy controls completed the task. Statistical analyses involving this test were thus performed on 23 IWA and 23 healthy controls.

#### 2.2.2 Phoneme identification task

The phoneme identification task assesses how consistently speech sounds (here /bA/-/dA/) are identified. We used the same task and stimuli as employed in Vandermosten et al. [31]. The task was presented as a two-alternative forced choice identification task. Participants were instructed to decide whether the stimulus they heard sounded more like a /bA/ or more like a /dA/. The stimuli were created based on a naturally spoken /bA/. The first 100 ms of the second formant (F2) of this syllable was linearly interpolated in 10 steps to create the stimuli, using Praat (Praat [46]; see Vandermosten et al. [31] for more details). The difference between /bA/ and /dA/ solely relies on the F2 slope, this way a gradual continuum was created between these speech sounds (fig. 1C). Thus, distinguishing between the two speech sounds relies mostly on dynamic cues, namely the discrimination of the spectral changes of F2 over time (i.e., whether the F2 slope is rising or falling). During the task, each of the 10 stimulus steps was presented 8 times in a randomized order, i.e., 80 trials. At the start of the task, the two speech sounds at the extremities of the stimulus spectrum were presented as reference practice trials. The stimuli were calibrated and presented monaurally at 70 dB SPL. The software APEX was used to present the task [44].

The amount of /dA/ responses for each stimulus step was taken and divided by 8 (i.e., number of presentations per stimulus step), to arrive at the proportion of /dA/ responses. This allowed us to fit a psychometric curve on the data points using the toolbox Psignifit in MATLAB (https://github.com/wichmann-lab/psignifit) [43]. This toolbox allows to fit subject-specific guess and lapse rates, thereby we avoided making assumptions about performance at the extremities of the stimulus continuum, hence the slope was not affected by such assumptions. As borders for the guess rate, we defined a range between 0 and 0.89 and for the lapse rate a range between 0 and 0.1 on the scale of proportion of /dA/ responses. We used uniformly distributed priors in order to avoid biasing the definition of the lapse and guess rate. Figure 2C shows the psychometric curves averaged by group. Subsequently, the slope at the subject-individual 50% point was computed using the function getSlope from the same toolbox and was used for statistical analyses. It is an indicator of how consistently participants were able to categorize the stimulus steps, which is indicated by the steepness of the slope.

In the aphasia group, 26/29 IWA completed the phoneme identification task, while 3 IWA experienced the task as too difficult after some initial trials. All 23 healthy controls completed the task. As a quality check, the confidence intervals of the lapse and guess rates were analyzed. Participants whose confidence interval of either of the asymptotes included 0.5 on the y-axis (i.e., the proportion of /dA/ responses; fig. 2C) were excluded from the analysis, i.e., 8 IWA and 4 healthy controls. Thus, all statistical analyses involving this test were performed on 18 IWA and 19 healthy controls.

#### 2.2.3 Phonological higher-level tasks

In order to answer the second research question, i.e., the potential link between acoustic-phonemic tasks and phonological processing, we used two tests, i.e., phonological word fluency and the phonology subtest of the ScreeLing. We used these measures to check whether scores on the RTD and phoneme identification task would predict the performance of IWA on these phonological higher-level tasks (within aphasia group analysis).

##### Phonological word fluency

We administered the phonological word fluency subscale of the CAT-NL [47]. Participants were required to enunciate as many words as possible that start with the letter ‘s’ within one minute. The score consisted of the number of correct words expressed. Phonological word fluency tasks require recruitment of linguistic functions, such as phonological processing and knowledge. However, note that phonological fluency tasks also involve cognitive functions, such as attention, executive functions and memory [48, 49, 50, 51]. We therefore controlled the linear models for the influence of cognitive functions, namely attention, executive functions and memory, in addition to controlling for hearing function.

##### Phonology subtest of the ScreeLing

The phonology scale consists of four tasks, i.e., spoken word repetition, reading out loud, minimal pair discrimination and initial phoneme identification. The two first tasks of the phonology subtest require the participant to produce speech as an answer, whereas the latter two tasks are receptive, i.e., the participant has to point at the answer. Specifically, the minimal pair discrimination task was auditorily presentated, i.e., 2 words followed by “were the two words you heard identical?”, and participants could point to yes or no or say it out loud. The initial phoneme identification task was presented simultaneously visually and auditorily, i.e., “what is the first letter of ‘word’?”, in reply to which participants got 4 multiple choice options and they could point at the answer or say it. Each of the tasks consists of 6 items, hence the total score on the phonology subtest is 24.

#### 2.2.4 Nuisance variables

We used measures of hearing and cognitive functioning as nuisance variables to take into account their potential influence on the dependent variable in the statistical models (section 2.3).

##### Hearing

Hearing thresholds were assessed via pure tone audiometry (PTA) at frequencies ranging from .25 to 4 kHz. The Fletcher index (average of thresholds at 1, 2 and 4 kHz) was calculated per ear and subsequently averaged across both ears. The thresholds did not differ between IWA and healthy controls (t= 0.582, df= 49.499, p= 0.563).

##### Cognitive functions

We administered the Oxford Cognitive Screen-NL (OCS) as cognitive test [52]. This validated test was designed to be language-independent, such that cognitive functioning can be disentangled from language functioning. Here, we used the subscales attention (i.e., crossing out target shapes among distractor shapes), executive functions (i.e., connecting circles and triangles in alternation in descending order of size) and memory (i.e., free recall and recognition of words and shapes) to calculate a composite score of cognition. The aphasia group had significantly lower cognitive scores than the healthy control group (t= −4.905, df= 33.759, p= <0.001).

### 2.3 Statistical analyses

Statistical analyses were performed in R [53]. We used parametric tests and then checked whether the normality assumptions were met (supplementary table S.2). If this was not the case, we conducted and reported non-parametric tests.

#### Research question 1

A two-tailed, unpaired Student’s t-test was performed to analyze group differences on the RTD and a two-tailed, unpaired Wilcoxon test to analyze differences on the phoneme identification tasks between the aphasia group and the healthy control group. The scores on the RTD task were log-transformed for statistical analyses because the outcome scores were logarithmically distributed, which was expected given the nature of the stimuli. Both the RTD and the phoneme identification task were auditorily presented and thus, require functional peripheral hearing. Older adults, independent of having aphasia or not, are prone to age-related hearing loss (i.e., presbyacusis) [54]. To account for individual differences in age-related hearing loss across all participants, we statistically controlled for its influence in a second step of the group comparison analysis. Furthermore, the two tasks also involve cognitive processes, such as attention and executive functioning. IWA frequently show concomitant cognitive impairments and thus, we also statistically controlled for the influence of cognitive functioning differences. Thus, to see whether the group effect would uphold when controlling for these variables, we added hearing levels (i.e., the Fletcher index) and cognition (i.e., composite score of OCS subtests of attention, executive functions and memory) to the models. For the linear model used for the RTD task, we used the following syntax: *task scores ~ group + hearing + cognition*. For the phoneme identification task we used a generalized additive model. We first checked which independent variables used more than one base function (i.e., is non-linear), which was hearing, and then applied the following syntax: *task scores ~ group + s(hearing) + cognition*.

An individual deviance analysis, as described in previous literature [13, 38, 55], was performed on the RTD and phoneme identification task. In essence, for this analysis a reference distribution was created based on a trimmed control group, and IWA were considered to deviate from this norm when their score exceeded 1.65 SD. More specifically, in a first step the lowest performing 5% of the control group were removed from the control group, which will be referred to as trimmed control sample. The mean and SD of the trimmed control sample were then used to standardize the raw task scores of all participants (IWA and all healthy controls). The deviance threshold was then defined at 1.65 SD for the RTD task and for the phoneme identification task at −1.65 SD of the z-scored distribution. Scores below the deviance threshold were viewed as deviant from the control sample (see supplementary material for more details on the implementation).

#### Research question 2

To investigate whether performance on the acoustic and phonemic tasks would predict performance at phonological processing tasks in IWA, we employed a linear model with the following syntax: *phonological task scores ~ acoustic-phonemic task scores + hearing + cognition*. We again controlled for the influence of hearing and cognition. An ANOVA was performed to test predictors of these linear models. The p-values were corrected for multiple comparisons (n=2, i.e., 2 phonology-level tests) using the false discovery rate (FDR) method [56].

## 3 Results

### 3.1 Research question 1: Comparison of the aphasia group with the healthy control group

We compared the RTD threshold of the aphasia group to the healthy control group and found a significant group difference (p = 0.001) (fig. 2B). All results are shown in table 2. IWA displayed on average a larger RTD threshold than healthy controls, meaning that they needed larger differences in rise time between the reference stimulus (rise time of 15 ms) and the deviant stimuli for discrimination. The group difference remained significant even after controlling for hearing and cognition (p= <0.001) (table 2).

**Table 2:**
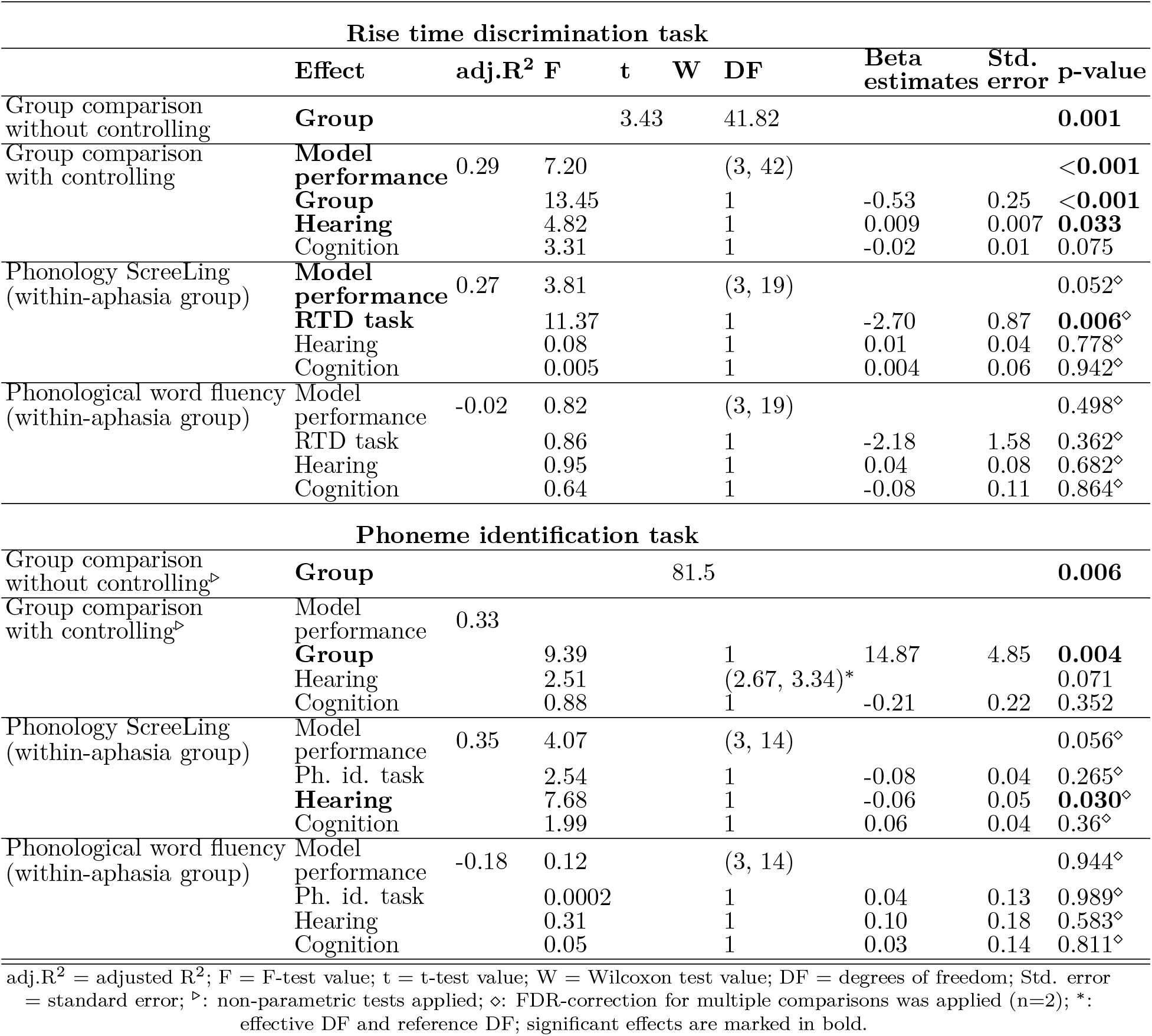
Results.

We also compared the phoneme identification slopes of the aphasia group to the healthy control group. We found a significant group difference (p = 0.006) (fig. 2D, table 2). IWA displayed on average less steep slopes of the psychometric function fitted to their data than healthy controls (fig. 2C), meaning that they did not classify the speech sounds as consistently in the same category as healthy controls. The group difference remained significant when controlling for inter-individual variability in hearing and cognition (p= 0.004). Neither hearing nor cognition significantly contributed to the model (table 2).

#### 3.1.1 Individual deviance analysis

We analyzed whether each IWA was deviant from the control group on the RTD and phoneme identification task. The original control group (n=23) was trimmed by removing two controls for the RTD task and one for the phoneme identification task, resulting in trimmed control samples of 21 and 22 participants respectively (see supplementary material for more details). After standardization of the scores based on the trimmed control group, we found that 12 out of 23 (52.2%) IWA were deviant from healthy controls on the RTD task. 10 out of 18 (55.5%) IWA were deviant from healthy controls on the phoneme identification task (supplementary table S.3). The groups’ distributions after standardisation of the scores relative to the deviance threshold are visualized in supplementary figure S.3. Taking the two tasks together, in total 19 out of 25 IWA (76%) were deviant on at least one of the tasks, meaning that three quarters of the aphasia sample had an impairment on at least one of the two auditory lower-level processing tasks.

### 3.2 Research question 2: Relation between acoustic-phonemic processing performance and phonological processing

We investigated whether RTD scores would predict the outcomes on the phonology subtest of the ScreeLing and the phonological word fluency test within the aphasia group. We found that the RTD thresholds significantly predicted scores on the phonology subtest of the ScreeLing (p= 0.006) (table 2 and figure 3). The larger the RTD thresholds were in IWA (i.e., the lower the performance), the lower the score was on the phonology subtest. This effect was present even though we controlled for inter-individual variability in hearing and cognitive functioning. Performance on the RTD task did not predict outcomes on the phonological word fluency task (p= 0.362). None of the factors we controlled for in the statistical model had a significant contribution to either of the models. Thus, the scores on the phonology ScreeLing subtest were predicted by the RTD scores above and beyond individual variations in hearing or cognitive functioning.

**Figure 3:**
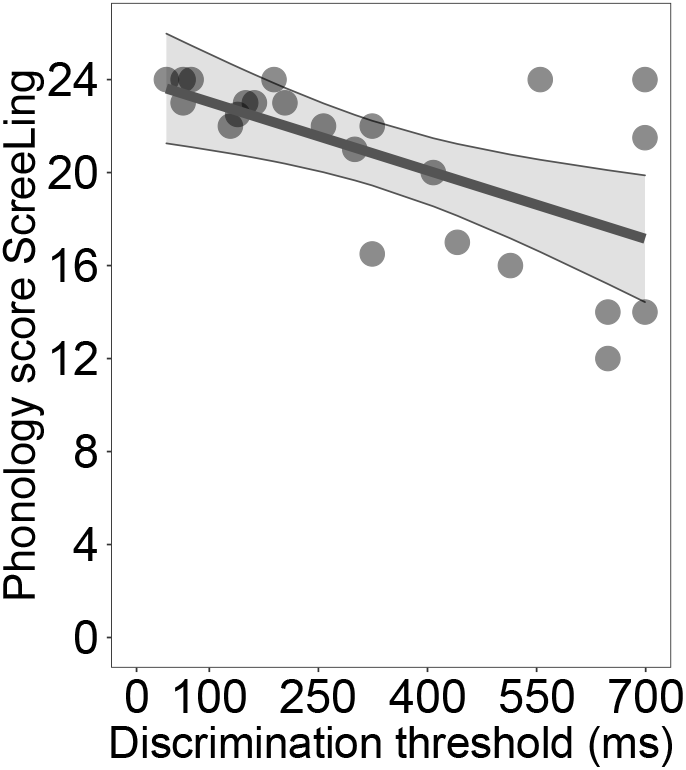
Performance on the RTD task predicts performance on the phonology subtest of the ScreeLing within IWA. **IWA** that detected differences in rise times only when the difference was relatively larger between standard and deviant stimuli (=lower performance) also scored lower on the phonology subtest of the ScreeLing, which consists of a composite score of four tasks, namely spoken word repetition, reading out loud, minimal pair discrimination and initial phoneme identification.

We also analyzed whether performance on the phoneme identification task would predict performance on the phonological processing tests within the aphasia group. We found that the phoneme identification slopes did neither predict scores on the phonology subtest of the ScreeLing (p=0.265) nor on the phonological word fluency test (p=0.989) within the aphasia group (table 2).

## 4 Discussion

We investigated acoustic and phonemic processing in individuals with post-stroke aphasia and age-matched healthy controls. Specifically, we administered two auditory tasks that rely on dynamic amplitude and spectral changes, which have not been investigated before in aphasia. The RTD task requires participants to detect small changes in amplitude. The phoneme identification task measures how consistently spectro-temporally varying, ambiguous speech sounds are classified in the same phonemic category and reflects how robust phoneme representations are (fig. 1). Assessing these tasks in IWA thus allows us to gain knowledge about lower-level auditory processing mechanisms that are currently neither assessed in clinical practice, nor treated in therapy. Here we did find group differences on both tasks, demonstrating lower performance in the aphasia group in rise time processing and phoneme identification than in the control group, even after controlling for the influence of hearing levels and cognitive functioning (fig. 2, table 2). We also observed that more than half of IWA were deviant from the control group on the RTD and on the phoneme identification task (supplementary fig. S.3). Taken both tasks together, three quarters of IWA were impaired at least at one of the tasks, and thus had an impairment at auditory spectro-temporal processing (supplementary table S.3).

Given that dyslexia research has found evidence for cascading effects from lower-level to higher-level processing impairments and aphasia research a link between auditory and phonological processing, we investigated whether performance of IWA on the acoustic and phonemic processing tasks can predict performance at two phonological processing tests, namely phonological word fluency and the phonology subtest of the ScreeLing (consisting of four tasks, i.e., spoken word repetition, reading out loud, minimal pair discrimination and initial phoneme identification). Indeed, we found that performance on the RTD task predicted performance on the phonology subtest of the ScreeLing, revealing that IWA who displayed lower performance on the RTD task also had lower scores on the phonology subtest of the ScreeLing (fig. 3, table 2). However, performance on the phoneme identification task did not significantly predict scores at any of the phonological tests. We will discuss the implications of these findings in detail here below.

In the current study, dynamic acoustic processing, measured via the RTD task, was lower in IWA than in healthy controls. Rise time processing has never been tested before in IWA, but our results are in line with previous studies investigating acoustic processing via non-dynamic psychoacoustic experiments [7, 25, 26, 27]. However, Oganian and Chang [12] demonstrated that processing of amplitude envelope modulations at the onset of speech sounds, i.e., acoustic onset edges and their slope steepness, rather than the absolute amplitude (i.e., static), is a crucial cue for speech comprehension. Therefore, assessing the dynamic processing of the rise time in IWA provides more specific information about potential impairments in speech envelope processing than static amplitude discrimination tasks. Deficient processing of the rise time during real-life speech can have adverse effects on understanding speech, because of its contribution to parsing the continuous speech stream into sublexcial and lexical segments [20]. Moreover, given that syllable stress influences the rise time [12, 19, 20], deficient rise time processing may lead to impaired processing of the syllable stress and thus, it becomes harder to follow the speech prosody and to comprehend speech in an efficient way.

Here, we found that more than half of IWA had impaired rise time processing, hence demonstrating that the assessment of lower-level auditory processing in IWA is important. As a matter of fact, 6 of our participants with aphasia were not able to perform this task because it was too difficult, potentially exhibiting an even larger proportion of IWA to be impaired on this task. We did not inquire about the specific reasons why these participants refused to proceed with the task. This would be important to explore in the future, should the task be considered for diagnostics. Overall, we suggest that being aware of an auditory processing impairment in a patient with aphasia could be useful for setting up an intervention plan targeting rise time processing and for following up on the recovery progress.

Execution of the RTD task also requires functioning sensory hearing and cognitive processes. Therefore, we statistically controlled for the variance explained by these factors in our analyses. The group effect remained significant. Nonetheless, we acknowledge that the administration of the RTD task would be difficult or impossible in patients with more severe conditions of cognitive impairment, motor impairments or hemineglect, which often occur in the acute phase after stroke, or in patients with severe hearing loss, thereby presenting a limitation for using the RTD task for individual diagnostics. This is also true for the phoneme identification task.

For efficient speech comprehension, certain cues of within- and between-speaker variability need to be inhibited in order to correctly identify phonemes [30]. The speech sounds used in the current phoneme identification task were artificially created to vary at different levels of ambiguity between /bA/ and /dA/, with the only difference between the sounds relying on spectro-temporal changes of the second formant within the first 100 ms after onset. In order to define the inter-category boundary between these phonemes and to consistently classify the same ambiguous stimuli into the same phoneme category, finetuned auditory spectro-temporal processing skills are essential. Thus, multiple processes are necessary for this task, i.e., sensitive auditory spectro-temporal processing, neglecting the variance within-speech sound category and linking the speech sound to phoneme representations in the brain, with the latter two processes being dependent on the first one.

Our results revealed that IWA identify phonemes less consistently than healthy, age-matched controls. Thus, it seems that IWA at group-level have less robust phoneme representations. Same as for the RTD task, the group difference remained significant after controlling for the variance explained by hearing and cognitive functioning. The group difference result is in line with Fink et al. [7], who found decreased performance in IWA on a word-level phoneme discrimination task that relied on temporal cues (i.e., voice onset time difference). The task by Fink et al. [7] did however not take into account dynamic spectral changes, which are especially important for processing formant transitions. Robson et al. [8] did explore dynamic spectro-temporal modulations of sounds and also found decreased performance in individuals with severe Wernicke’s aphasia. Besides the difference in nature of stimuli, the current study also expands the finding of Robson et al. [8] to a more broadly recruited group of IWA, whose severity and type of aphasia is more heterogeneous.

An impairment in consistently identifying phonemes may be due to inefficient auditory spectro-temporal processing, but could also be impacted by difficulties with neglecting within-phoneme category variance or with linking the sound to the correct phoneme representation. Administering a phoneme discrimination task in addition to the phoneme identification task may be useful to disentangle these processes in the future. The phoneme discrimination task requires participants to indicate whether two speech sounds are the same (see supplementary experiment of Schevenels et al. [57]). Hence, categorizing phonemes is not necessary and a potential impairment on this task would be due to poor spectro-temporal processing, eliminating the possible influence of other processes. In future studies, we suggest to administer a phoneme discrimination task as a complement to the phoneme identification task to isolate the involved processes.

Not only did we find lower performance on the phoneme identification task in IWA, but we also detected that more than half of IWA have an impaired performance on this task, as evidenced by the individual deviance analysis. If we take into account the participants that had to be excluded from the analysis of the phoneme identification task (n=8 IWA) because of too poor performance to fit a meaningful psychometric function, then 18 out of 26 IWA (69.23%) were deviant from healthy controls on the phoneme identification task. The large proportion of IWA deviant on this task shows that it is important to assess phonemic processing in aphasia and train phonemic representations during the recovery process. This result is in line with findings of Robson et al. [8], who also reported relatively high proportions of deviance in IWA on 3 dynamic modulation tasks.

Taking a look at the overlap of deviance between the RTD task and the phoneme identification task in IWA, we saw that only a limited amount of them showed concordant deviance. In fact, 62.5% of IWA did not show an overlap of deviance between tasks, meaning that they were deviant on the RTD task but not on the phoneme identification task or vice versa. Even though both tasks require analysis of low-level auditory aspects, this shows that the two tasks do, at least partially, not measure one same construct of auditory spectro-temporal processing. While the RTD task assesses dynamic changes in amplitude, the phoneme identification task measures dynamic changes in frequency. Thus, it is possible that some IWA have more difficulties with processing amplitude changes, whereas others struggle more with dynamic changes in frequency. Still others might face difficulties in processing both aspects. However, given the small sample size in this study, we cannot draw strong conclusions. Nonetheless, it would be interesting to further investigate this in the future.

An alternative explanation for the limited overlap of deviance between tasks could be the task-specific cognitive processes involved. For both tasks, auditory attention needs to be allocated to the stimuli and stimuli need to be analyzed in order to take a decision, drawing on executive functions. However, the RTD task may require more involvement of short-term memory because three sounds have to be kept in short-term memory in order to compare them to each other to take a decision. The phoneme identification task on the other hand requires participants to take an immediate decision after listening to one sound per trial, which does not require short-term memory involvement. Participants might however compare the sounds across trials and thus, the phoneme identification task may also draw on short-term memory, yet less intensively. IWA may or may not have impaired short-term memory and if so, to varying extents. Hence, we cannot rule out that the difference in short-term memory involvement in the tasks may partly explain the limited overlap in deviance between the RTD and the phoneme identification task.

We also explored whether auditory processing would predict phonological processing in IWA. In the past, theoretical models of speech processing have viewed the different steps to be sequential and unidirectional, i.e., the auditory phonological analysis is followed by integration into the phonological lexicon, which is in turn followed by activation of the semantic system [58, 59]. More recent models, however, show that different speech processing levels interact bidirectionally with each other[35, 36]. In both models, an auditory processing impairment may propagate onto higher-level processes. This hypothesis has been tested in individuals with developmental dyslexia, where evidence has shown that rise time processing predicts phonological processing performance and literacy measures [13, 14, 15, 16, 17]. Similarly, we found that performance on the RTD task predicts higher-level phonological processing in IWA, as measured by the phonology subtest of the ScreeLing. This test analyzes phonological processing at a metalinguistic level, e.g., phonological awareness. Using a different type of spectro-temporal processing task, Robson et al. [8] reported an association between auditory processing and the phonological discrimination task, which is thus in line with the current results.

Integrating our findings, we established that more than half of IWA showed impaired performance on the RTD task and this performance relates to phonological processing. This could have interesting implications for diagnosis and therapy of aphasia. Could therapy targeting improvement of rise time processing possibly show transfer effects on phonological processing in IWA? Taking a look at intervention research, a study in children at cognitive risk for dyslexia has shown that an intervention with enhanced envelope listening improved RTD performance [60]. In IWA, Szymaszek et al. [61] showed that a training in temporal processing improved not only temporal processing performance, but also transferred to language comprehension tasks. Hence, having tools available to assess auditory processing in IWA paves the way for developing according treatment methods in the future, which may even show transfer effects to higher-level language processing.

In contrast to the RTD task, we found that the phoneme identification task did not predict performance at either of the phonological tests in IWA. This stands in opposition to our hypotheses that were based on two studies. First, Robson et al. [8] found a link between auditory and phonological processing in IWA, although in a small sample size and with a different auditory task. Second, in dyslexia research, the phoneme identification task has been linked to higher-level phonological processing [18], suggesting that deficits at the phonemic processing level do propagate onto higher-level speech processing mechanisms in dyslexia. The lack of such a result here might be due to a small sample size (n=18, whereas n=23 for RTD task). Studies with a larger sample size may shine light on this in the future.

Unlike the performance of IWA on the ScreeLing phonology subtest, their performance on the phonological word fluency test was not predicted by the rise time processing performance. We have two possible explanations for this result. First, we suggest that the amount and intensity of cognitive involvement is larger during the word fluency test than for the ScreeLing phonology subtest. The phonological word fluency task is time-constrained, attention-heavy and participants need to make use of executive functions, such as strategy formation, verbal memory (word retrieval), word knowledge and giving goal-directed responses according to the task rule [48, 49, 50]. Phonological word fluency is not solely used as a task to measure language performance, but also to measure executive dysfunction [49, 50]. The ScreeLing phonology subtest on the other hand contains tasks requiring auditory attention, verbal shortterm memory and decision-making processes. Thus, comparing the cognitive processes involved in these two tests, the word fluency test involves more and more costly mechanisms than the phonology subtest of the ScreeLing. Second, the cognitive processes involved in the RTD task, as well as the phoneme identification task, are more similar to the ones involved in the ScreeLing phonology subtest than to those involved in the word fluency test. As discussed earlier, the acoustic and phonemic tasks involve attention, short-term memory (to varying extents) and decision-making. These cognitive processes are similar to the ones involved in the ScreeLing phonology subtest, but differ from those involved in the word fluency task.

In conclusion, our results show that the RTD task and the phoneme identification task were each able to distinguish IWA from healthy controls. Moreover, we found that three-quarters of our aphasia sample do suffer from acoustic or phonemic processing problems, in addition to potential higher-level language processing impairments. Assessment of auditory processing is however currently not done in the clinic when it comes to diagnosing aphasia. Additionally, we demonstrated that performance on the RTD task predicted phonological processing skills. Future development of norm scores of the acoustic and phonemic tasks would allow to formally diagnose auditory spectro-temporal processing impairments in IWA and would thus help SLPs to target therapy towards those aspects. Both the acoustic and phonemic task only require a tablet and headphones and would thus be relatively easy to implement in the clinical context (hospitals, rehabilitation centra or SLP practices). Patients with aphasia in the acute, subacute and chronic phase after stroke that are testable for language tests are also able to perform these tasks, which do not require a verbal response from patients. Both tasks take between 5 and 10 minutes administration time and display the results immediately after completion. Due to their efficiency and feasibility, the RTD and the phoneme identification tasks may be useful for diagnosis and follow-up of aphasia in the clinical context. Further research would be required in order to validate the tasks and develop norm scores for both tasks.

## Data Availability

After completion of the review process the preprocessed data will be made available.

## Funding Sources

Research of Jill Kries was supported by the Luxembourg National Research Fund (FNR) (AFR-PhD project reference 13513810). Pieter De Clercq was financially supported by the Flanders Wetenschappelijk Onderzoek (FWO) SB grant, No. 1S40122N. Robin Lemmens is a senior clinical investigator supported by the FWO. The presented study also received funding from the European Research Council (ERC) under the European Union’s Horizon 2020 research and innovation programme (Tom Francart; grant agreement No. 637424). Furthermore, this study was financially supported by the FWO grant No. G0D8520N.

## Additional information

No conflicts of interest, financial or otherwise, are declared by the authors.

## Acknowledgements

The authors would like to thank all participants, especially all the brave participants with aphasia and their partners, family or friends that support them. Furthermore, the authors would like to thank Dr. Klara Schevenels for helping with recruitment of participants with aphasia and with setting up the auditory-phonemic tasks. We are also grateful for the help of Dr. Benjamin Dieudonné, Dr. Toivo Glatz and Dr. Jonas Vanthornhout with methodological implementations. A big thanks goes to everyone who helped with data collection and recruitment: Janne Segers, Rosanne Partoens, Charlotte Rommel, Dr. Ramtin Mehraram, Ines Robberechts, Laura Van Den Bergh, Anke Heremans, Frauke De Vis, Mouna Vanlommel, Naomi Pollet, Kaat Schroeven, Pia Reynaert and Merel Dillen.

## Author contributions statement

MV and JK conceived and designed research; JK set up the protocol and tasks; JK and PDC collected data; JK processed and analyzed data; JK and MV interpreted results of experiments; JK prepared figures; JK drafted manuscript; MV, JK, TF, RL and PDC edited and revised manuscript; JK, PDC, RL, TF and MV approved final version of manuscript.

## Supplementary material

### Recruitment flowchart

**Figure S.1:**
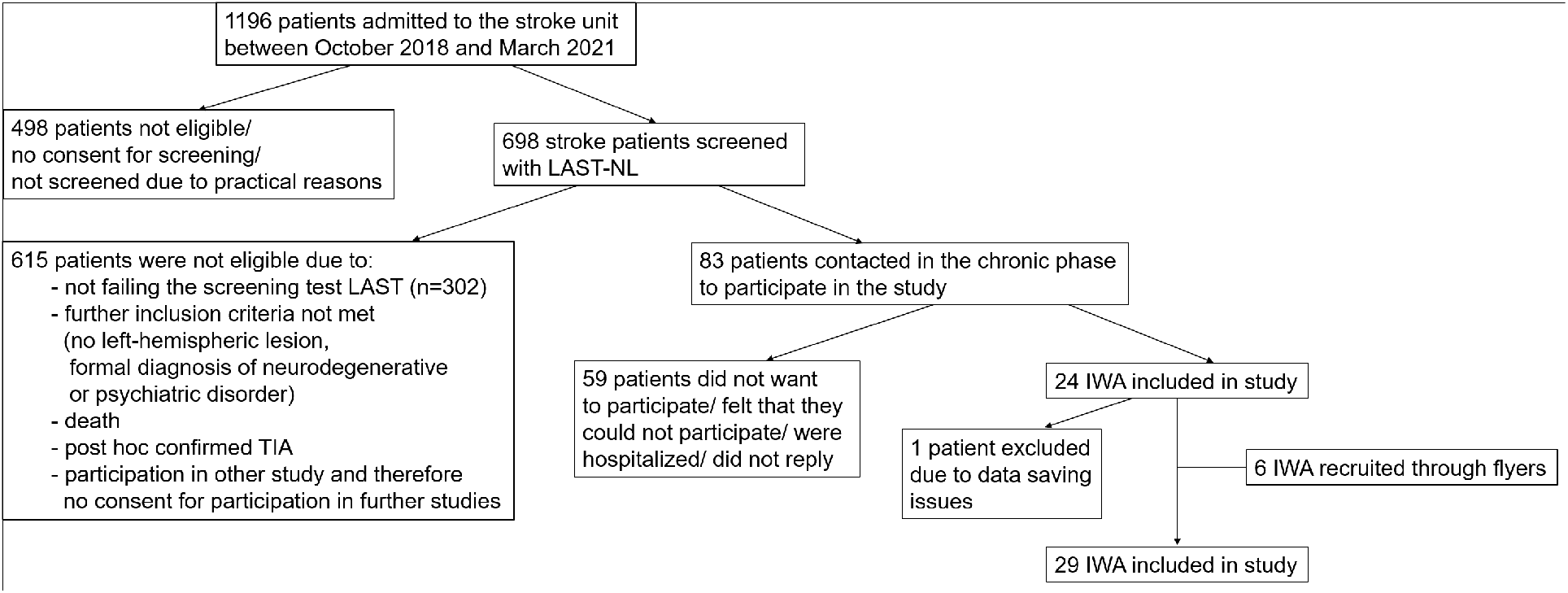
Flowchart of the recruitment procedure of individuals with aphasia as described in section 2.1 of the paper.

### Visualization of demographic and diagnostic variables

**Figure S.2:**
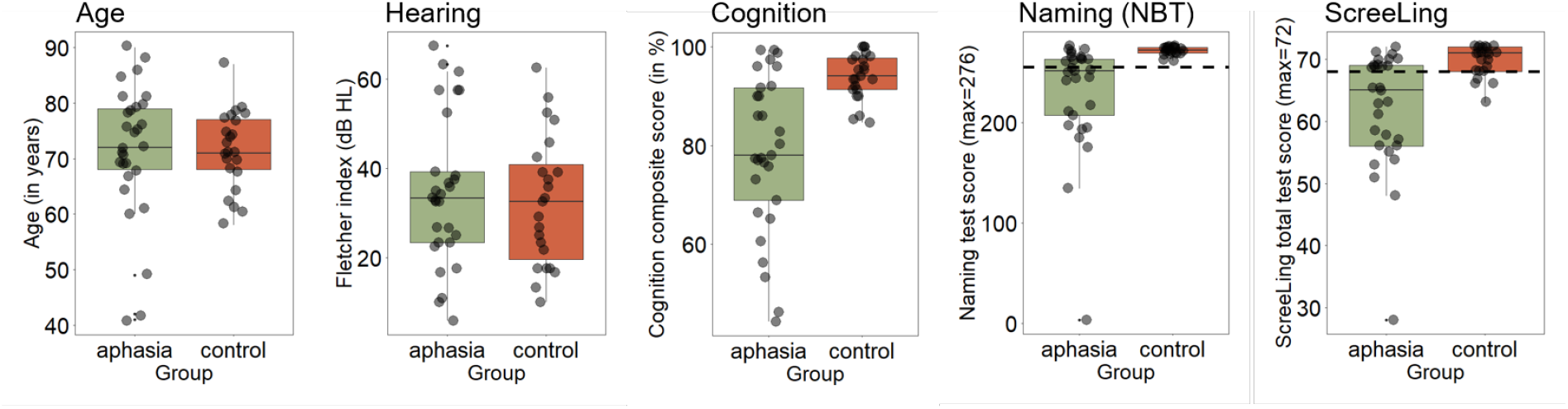
This figure shows demographic and diagnostic variables by group of the variables age, hearing, cognition, naming test and diagnostic language test. The dashed lines on the two right most figures correspond to the cut-off threshold of those tests.

### Education levels

**Table S.1:** Contingency table for education levels per group.

| Education level<br>(years of education) | Aphasia group<br>n (%) | Control group<br>n (%) |
| --- | --- | --- |
| Primary school<br>(6 years) | 2 (6.90%) | 0 (0.00%) |
| Secondary school<br>(12 years) | 10 (34.48%) | 5 (21.74%) |
| College degree<br>(15 years) | 8 (27.59%) | 8 (34.78%) |
| University degree<br>(17 years) | 9 (31.03%) | 8 (34.78%) |
| Doctoral degree<br>(21 years) | 0 (0.00%) | 2 (8.70%) |

### Normality assumptions of variables used for statistical analyses

**Table S.2:** Shapiro-Wilk and Levene’s test results to check the normality assumptions.

| Effect | Test | W-value | F-Value | DF | p-value |
| --- | --- | --- | --- | --- | --- |
| **Rise time discrimination task** |  |  |  |  |  |
| Group comparison<br>without controlling | Normality (Shapiro-Wilk test) | 0.96 |  |  | 0.205 |
|  | Homoscedasticity (Levene's test) |  | 2.03 | 1, 44 | 0.160 |
| Group comparison<br>with controlling | Normality (Shapiro-Wilk test) | 0.97 |  |  | 0.332 |
| Phonology ScreeLing<br>(within aphasia group) | Normality (Shapiro-Wilk test) | 0.97 |  |  | 0.882 |
| Phonological word fluency<br>(within aphasia group) | Normality (Shapiro-Wilk test) | 0.92 |  |  | 0.094 |
| **Phoneme identification task** |  |  |  |  |  |
| Group comparison<br>without controlling | **Normality (Shapiro-Wilk test)** | 0.93 |  |  | **0.031** |
|  | Homoscedasticity (Levene's test) |  | 2.33 | 1, 35 | 0.135 |
| Group comparison<br>with controlling | **Normality (Shapiro-Wilk test)** | 0.91 |  |  | **0.01** |
| Phonology ScreeLing<br>(within aphasia group) | Normality (Shapiro-Wilk test) | 0.96 |  |  | 0.658 |
| Phonological word fluency<br>(within aphasia group) | Normality (Shapiro-Wilk test) | 0.96 |  |  | 0.694 |

DF = degrees of freedom; significant effects are marked in bold, meaning that the data are not meeting the normality assumption.

### Individual deviance analysis

The individual deviance analysis allows to see which individuals with aphasia are deviant from the control group on the RTD task and the phoneme identification task (see paper for more details on the method). For the RTD task, the control sample was normally distributed after removing the lowest performing 5% (≥ percentile 95) of the control group (Shapiro-Wilk normality test: W = 0.92, p-value = 0.122). For the phoneme identification task, the control sample was also normally distributed after removing the lowest performing 5% (≤ percentile 5) of the control group (Shapiro-Wilk normality test: W = 0.95, p-value = 0.517). For the threshold estimation, all participant scores were standardized by subtracting the mean of the trimmed control sample and then dividing by the SD of the trimmed control sample. The deviance threshold was then defined at −1.65 SD (for the phoneme identification task) or 1.65 SD (for the RTD task) of the z-scored distribution.

**Figure S.3:**
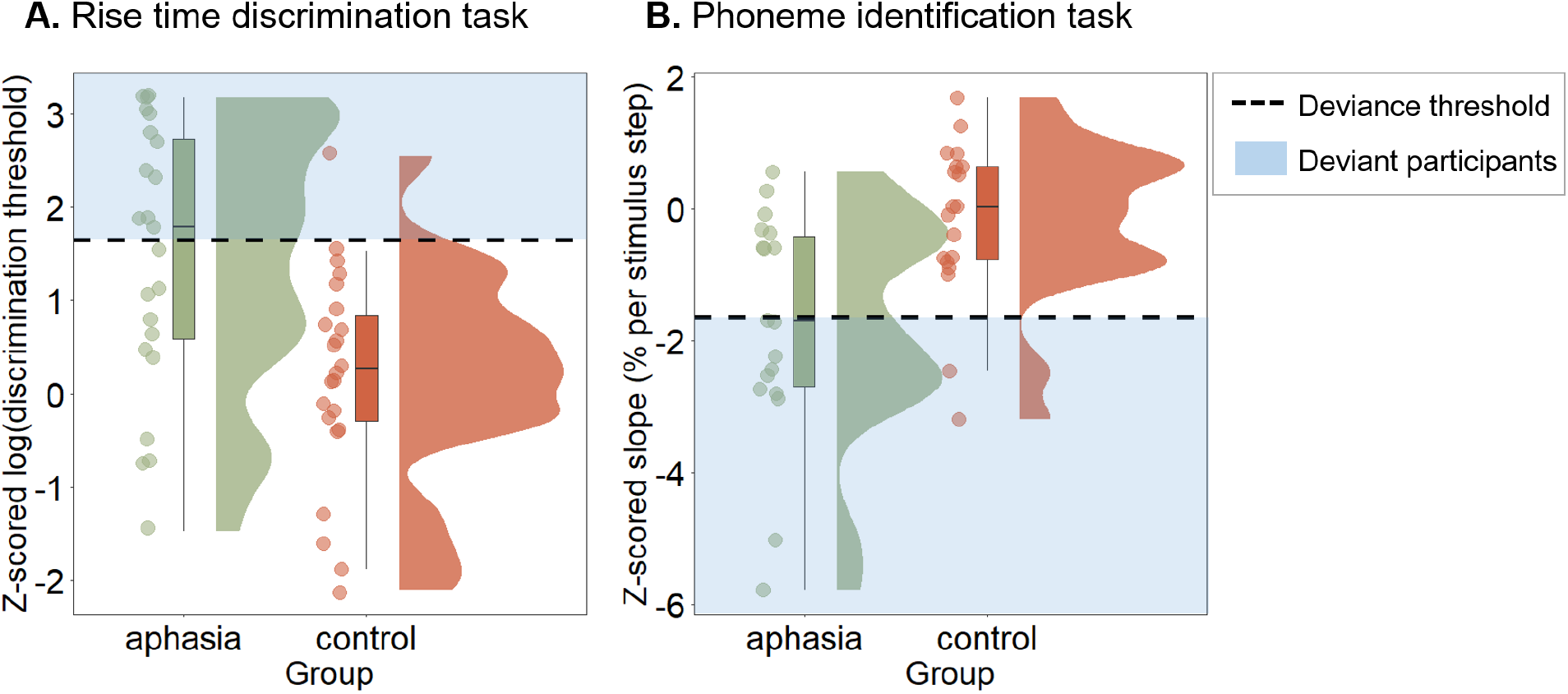
Individual deviance analysis of the acoustic and phonemic processing tasks. **A.** Visualization of the deviant participants for the RTD task. **B.** Visualization of the deviant participants for the phoneme identification task.

**Table S.3:** Number of participants and percentage of deviance on the acoustic and phonemic tasks.

|  | Aphasia group<br>n(%) | Control group<br>n(%) |
| --- | --- | --- |
| **Rise time discrimination task** |  |  |
| deviant | 12 (52.17%) | 1 (4.35%) |
| not deviant | 11 (47.83%) | 22 (95.65%) |
| **Phoneme identification task** |  |  |
| deviant | 10 (55.56%) | 2 (10.53%) |
| not deviant | 8 (44.44%) | 17 (89.47%) |
| **Overlap in deviance between the 2 tasks** |  |  |
| overlap | 6 (37.50%) | 17 (89.47%) |
| no overlap | 10 (62.5%) | 2 (10.53%) |
| **Deviance on at least one of the 2 tasks** |  |  |
| yes | 19 (76%) | 3 (13.04%) |
| no | 6 (24%) | 20 (86.96%) |

## Notes

### Competing Interest Statement

The authors have declared no competing interest.

## References

[1] Pasley, B. N. & Knight, R. T. Decoding speech for understanding and treating aphasia. Progress in Brain Research 207, 435–456 (2013).

[2] Rohde, A. et al. Diagnosis of aphasia in stroke populations: A systematic review of language tests. PLoS ONE 13, 1–17 (2018).

[3] El Hachioui, H. et al. Screening tests for aphasia in patients with stroke: a systematic review. Journal of Neurology 264, 211–220 (2017).

[4] Royal College of Speech and Language Therapists. Clinical guidelines (2005). URL http://tcssexed.weebly.com/uploads/1/2/5/9/12593116/ebp_rcslt_clinical_guidelines.pdf.

[5] Visch-Brink, E., Links, P. & Hurkmans, J. Richtlijn linguïstische diagnostiek en therapie bij een verworven afasie, augustus 2012 (2012). URL https://klinischelinguistiek.nl/uploads/richtlijnlinguistischediagnost.pdf.

[6] American Speech-Language-Hearing Association. Preferred practice patterns for the profession of speech-language pathology (2004). URL www.asha.org/policy/.

[7] Fink, M., Churan, J. & Wittmann, M. Temporal processing and context dependency of phoneme discrimination in patients with aphasia. Brain and Language 98, 1–11 (2006).

[8] Robson, H., Grube, M., Lambon Ralph, M. A., Griffiths, T. D. & Sage, K. Fundamental deficits of auditory perception in wernicke’s aphasia. Cortex 49, 1808–1822 (2013). URL http://dx.doi.org/10.1016/j.cortex.2012.11.012.

[9] Shannon, R. V., Zeng, F.-G., Kamath, V., Wygonski, J. & Ekelid, M. Speech recognition with primarily temporal cues. Science 270, 303 (1995). URL https://www.proquest.com/scholarly-journals/speech-recognition-with-primarily-temporal-cues/docview/213563934/se-2.

[10] Zeng, F. G. et al. Speech recognition with amplitude and frequency modulations. Proceedings of the National Academy of Sciences of the United States of America 102, 2293–2298 (2005). URL www.pnas.orgcgidoi10.1073pnas.0406460102.

[11] Xu, L. & Pfingst, B. E. Spectral and temporal cues for speech recognition: Implications for auditory prostheses. Hearing Research 242, 132–140 (2008).

[12] Oganian, Y. & Chang, E. F. A speech envelope landmark for syllable encoding in human superior temporal gyrus. Science Advances 5, 1–13 (2019). URL https://www.science.org.

[13] Law, J. M., Vandermosten, M., Ghesquiere, P., Wouters, J. & De Bree, E. The relationship of phonological ability, speech perception, and auditory perception in adults with dyslexia. Frontiers in Human Neuroscience 8, 1–12 (2014). URL www.frontiersin.org.

[14] Richardson, U., Thomson, J. M., Scott, S. K. & Goswami, U. Auditory processing skills and phonological representation in dyslexic children. Dyslexia 10, 215–233 (2004).

[15] Pasquini, E. S., Corriveau, K. H. & Goswami, U. Scientific studies of reading auditory processing of amplitude envelope rise time in adults diagnosed with developmental dyslexia. Scientific Studies of Reading 11 (2007). URL https://doi.org/10.1080/10888430701344280.

[16] De Vos, A., Vanvooren, S., Vanderauwera, J., Ghesquière, P. & Wouters, J. A longitudinal study investigating neural processing of speech envelope modulation rates in children with (a family risk for) dyslexia. Cortex 93, 206–219 (2017). URL www.sciencedirect.comhttp://dx.doi.org/10.1016/j.cortex.2017.05.007.

[17] Vanvooren, S., Poelmans, H., De Vos, A., Ghesquière, P. & Wouters, J. Do prereaders’ auditory processing and speech perception predict later literacy? Research in Developmental Disabilities 70, 138–151 (2017). URL http://dx.doi.org/10.1016/j.ridd.2017.09.005.

[18] Boets, B. et al. Preschool impairments in auditory processing and speech perception uniquely predict future reading problems. Research in developmental disabilities 32, 560–570 (2011). URL https://pubmed.ncbi.nlm.nih.gov/21236633/.

[19] Hämäläinen, J. A., Rupp, A., Soltész, F., Szücs, D. & Goswami, U. Reduced phase locking to slow amplitude modulation in adults with dyslexia: An meg study. NeuroImage 59, 2952–2961 (2012).

[20] Goswami, U., Fosker, T., Huss, M., Mead, N. & Szucs, D. Rise time and formant transition duration in the discrimination of speech sounds: The ba-wa distinction in developmental dyslexia. Developmental Science 14, 34–43 (2011).

[21] Biedermann, F., Bungert, P., Dörrscheidt, G. J., Von Cramon, D. Y. & Rübsamen, R. Central auditory impairment in unilateral diencephalic and telencephalic lesions. Audiology and Neurotology 13, 123–144 (2008).

[22] Navarro-Orozco, D. & Sánchez-Manso, J. C. Neuroanatomy, Middle Cerebral Artery (StatPearls Publishing, Treasure Island, FL, US, 2022). URL https://www.ncbi.nlm.nih.gov/books/NBK526002/.

[23] Flowers, H. L. et al. Poststroke aphasia frequency, recovery, and outcomes: A systematic review and meta-analysis. Archives of Physical Medicine and Rehabilitation 97, 2188–2201 (2016).

[24] Hillis, A. E. Aphasia progress in the last quarter of a century. Neurology 69 (2007).

[25] Ilvonen, T. et al. The processing of speech and non-speech sounds in aphasic patients as reflected by the mismatch negativity (mmn). Neuroscience Letters 366, 235–240 (2004).

[26] Sidiropoulos, K., Ackermann, H., Wannke, M. & Hertrich, I. Temporal processing capabilities in repetition conduction aphasia. Brain and Cognition 73, 194–202 (2010). URL http://dx.doi.org/10.1016/j.bandc.2010.05.003.

[27] Stefanatos, G. A., Braitman, L. E. & Madigan, S. Fine grain temporal analysis in aphasia: Evidence from auditory gap detection. Neuropsychologia 45, 1127–1133 (2007). URL http://dx.doi.org/10.1016/j.neuropsychologia.2006.09.011.

[28] Hämäläinen, J. A., Salminen, H. K. & Leppänen, P. H. T. Basic auditory processing deficits in dyslexia: Systematic review of the behavioral and event-related potential/ field evidence. Journal of Learning Disabilities 46, 413–427 (2013). URL https://doi.org/10.1177/0022219411436213.

[29] Van Hirtum, T., Ghesquière, P. & Wouters, J. Atypical neural processing of rise time by adults with dyslexia. Cortex 113, 128–140 (2019).

[30] Binder, J. R. Phoneme perception. In Hickok, G. & Small, S. (eds.) Neurobiology of Language, chap. 37, 447–461 (Academic Press, London, UK, 2016).

[31] Vandermosten, M. et al. Adults with dyslexia are impaired in categorizing speech and nonspeech sounds on the basis of temporal cues. Proceedings of the National Academy of Sciences of the United States of America 107, 10389–10394 (2010). URL https://www.pnas.org/content/107/23/10389.

[32] Liebenthal, E., Binder, J. R., Spitzer, S. M., Possing, E. T. & Medler, D. A. Neural substrates of phonemic perception. Cerebral Cortex 15, 1621–1631 (2005).

[33] Desai, R., Liebenthal, E., Waldron, E. & Binder, J. R. Left Posterior Temporal Regions are Sensitive to Auditory Categorization. Journal of Cognitive Neuroscience 20, 1174–1188 (2008). URL https://doi.org/10.1162/jocn.2008.20081.

[34] Turkeltaub, P. E. & Coslett, H. B. Localization of sublexical speech perception components. Brain and Language 114, 1–15 (2010).

[35] Gwilliams, L., Linzen, T., Poeppel, D. & Marantz, A. In spoken word recognition, the future predicts the past. Journal of Neuroscience 38, 7585–7599 (2018). URL https://www.jneurosci.org/content/38/35/7585. https://www.jneurosci.org/content/38/35/7585.full.pdf.

[36] Gwilliams, L. & Davis, M. H. Extracting language content from speech sounds: the information theoretic approach. In Holt, L. L., Peelle, J. E., Coffin, A. B., Popper, A. N. & Fay, R. R. (eds.) Speech Perception, chap. 5, 113–139 (Springer Cham, Switzerland, 2022).

[37] Goswami, U. Sensory theories of developmental dyslexia: three challenges for research. Nature Reviews Neuroscience 16, 43–54 (2014). URL www.nature.com/reviews/neuro.

[38] Ramus, F. et al. Theories of developmental dyslexia: insights from a multiple case study of dyslexic adults. Brain 126, 841–65 (2003).

[39] Flamand-Roze, C. et al. Validation of a new language screening tool for patients with acute stroke. Stroke 42, 1224–1229 (2011). URL https://www.ahajournals.org/doi/abs/10.1161/STROKEAHA.110.609503. https://www.ahajournals.org/doi/pdf/10.1161/STROKEAHA.110.609503.

[40] Visch-Brink, E., Van de Sandt-Koenderman, M. & El Hachioui, H. ScreeLing (Bohn Stafleu Van Loghum, Houten, NL, 2010).

[41] Van Ewijk, E. et al. Nederlandse Benoem Test (Bohn Stafleu Van Loghum, Houten, NL, 2020).

[42] Anwyl-Irvine, A. L., Massonnié, J., Flitton, A., Kirkham, N. & Evershed, J. K. Gorilla in our midst: An online behavioral experiment builder. Behavior Research Methods 52, 388–407 (2020).

[43] MATLAB. version 9.1.0.441655 (R2016b) (The MathWorks Inc., Natick, Massachusetts, 2016).

[44] Francart, T., van Wieringen, A. & Wouters, J. APEX 3: a multi-purpose test platform for auditory psychophysical experiments. J Neurosci Methods 172, 283–293 (2008).

[45] Levitt, H. Transformed up-down methods in psychoacoustics. The Journal of the Acoustical Society of America 49 (1971).

[46] Boersma, P. & Weenink, D. Praat: doing phonetics by computer (2022). URL http://www.praat.org/.Computerprogram.

[47] Swinburn, K. et al. CAT-NL: Comprehensive Aphasia Test (Amsterdam: Pearson, 2004).

[48] Ruff, R. M., Light, R. H., Parker, S. B. & Levin, H. S. The psychological construct of word fluency. Brain and Language 57, 394–405 (1997).

[49] Henry, J. D. & Crawford, J. R. A meta-analytic review of verbal fluency performance following focal cortical lesions. Neuropsychology 18, 284–295 (2004).

[50] Sarno, M. T., Postman, W. A., Cho, Y. S. & Norman, R. G. Evolution of phonemic word fluency performance in post-stroke aphasia. Journal of Communication Disorders 38, 83–107 (2005).

[51] Schmidt, C. S. et al. Dissociating frontal and temporal correlates of phonological and semantic fluency in a large sample of left hemisphere stroke patients. NeuroImage: Clinical 23 (2019).

[52] Huygelier, H., Schraepen, B., Demeyere, N. & Gillebert, C. R. The Dutch version of the Oxford Cognitive Screen (OCS-NL): normative data and their association with age and socio-economic status. Aging, Neuropsychology, and Cognition 27, 765–786 (2019). URL https://doi.org/10.1080/13825585.2019.1680598.

[53] R Core Team. R: A Language and Environment for Statistical Computing. R Foundation for Statistical Computing, Vienna, Austria (2017). URL https://www.R-project.org/.

[54] Rankin, E., Newton, C., Parker, A. & Bruce, C. Hearing loss and auditory processing ability in people with aphasia. Aphasiology 28, 576–595 (2014). URL http://dx.doi.org/10.1080/02687038.2013.878452.

[55] Boets, B., Wouters, J., van Wieringen, A. & Ghesquière, P. Auditory processing, speech perception and phonological ability in pre-school children at high-risk for dyslexia: A longitudinal study of the auditory temporal processing theory. Neuropsychologia 45, 1608–1620 (2007).

[56] Benjamini, Y. & Hochberg, Y. Controlling the false discovery rate: A practical and powerful approach to multiple testing. Journal of the Royal Statistical Society 57, 289–300 (1995).

[57] Schevenels, K., Altvater-Mackensen, N., Zink, I., De Smedt, B. & Vandermosten, M. Aging effects and feasibility of statistical learning tasks across modalities. Aging, Neuropsychology, and Cognition 0, 1–30 (2021). URL https://doi.org/10.1080/13825585.2021.2007213. https://doi.org/10.1080/13825585.2021.2007213.

[58] Ellis, A. & Young, A. Human Cognitive Neuropsychology: A Textbook With Readings (1st ed.) (Psychology Press, London, 1996).

[59] Papathanasiou, I. & Coppens, P. Aphasia and Related Neurogenic Communication Disorders (Sudbury, MA: Jones and Bartlett, 2016).

[60] Van Herck, S. et al. Ahead of maturation: Enhanced speech envelope training boosts rise time discrimination in pre-readers at cognitive risk for dyslexia. Developmental Science 25, 1–12 (2022).

[61] Szymaszek, N., Wolak, T. & Szelag, E. The treatment based on temporal information processing reduces speech comprehension deficits in aphasic subjects. Frontiers in Aging Neuroscience 9, 1–11 (2017).

